# NMDAR mediated dynamic changes in m^6^A inversely correlates with neuronal translation

**DOI:** 10.1101/2022.06.01.494458

**Authors:** Naveen Kumar Chandappa Gowda, Bharti Nawalpuri, Sarayu Ramakrishna, Vishwaja Jhaveri, Ravi S Muddashetty

## Abstract

Epitranscriptome modifications are crucial in translation regulation and essential for maintaining cellular homeostasis. N6 methyladenosine (m^6^A) is one of the most abundant and well-conserved epitranscriptome modifications, which is known to play a pivotal role in diverse aspects of neuronal functions. However, the role of m^6^A modifications with respect to activity-mediated translation regulation and synaptic plasticity has not been studied. Here, we investigated the role of m^6^A modification in response to NMDAR stimulation. We have consistently observed that upon 5-minute NMDAR stimulation causes an increase in eEF2 phosphorylation. Correspondingly, NMDAR stimulation caused a significant increase in the m^6^A signal at 5 minutes time point, correlating with the global translation inhibition. The NMDAR induced increase in the m^6^A signal is accompanied by the redistribution of the m^6^A marked RNAs from translating to the non-translating pool of ribosomes. The increased m^6^A levels are well correlated with the reduced FTO levels observed on NMDAR stimulation. Additionally, we show that inhibition of FTO prevents NMDAR mediated changes in m6A levels. Overall, our results establish RNA-based molecular readout which corelates with the NMDAR-dependent translation regulation which helps in understanding changes in protein synthesis.

## Introduction

m^6^A is a widespread, abundant and well conserved internal RNA modification, which plays a pivotal role in regulating multitude of physiological and pathological processes. It is present on diverse classes of RNA molecules such as rRNAs, mRNAs, tRNAs, snRNAs, miRNAs, and long non-coding RNAs. In mammalian mRNA, m^6^A is primarily located near the stop codon, 3’ UTR, and long internal exons^1^. It is known to critically regulate several aspects of RNA metabolism, such as RNA stability, splicing, translation, translocation, localization, degradation, and transport ^2–4^. This modification is highly enriched in the brain and global levels are developmentally regulated ^5^.

In mammals, the m^6^A mark on RNA is dynamically and reversibly regulated by the action of RNA methyltransferases (writers) and demethylases (erasers)^6^. Methyl groups are added co-transcriptionally onto the adenosine nucleotide through a multicomponent methyltransferase complex. This complex consists of a catalytic core protein Methyltransferase-like 3 (METTL3), along with the adaptor proteins Methyltransferase-like 14 (METTL14) and Wilms tumour 1 associated protein (WTAP)^6^. This m^6^A mark can be reversed by the action of m^6^A demethylases, Fat mass and obesity-associated (FTO)^7^ and Alkbh5^8^. These demethylases primarily differ in their substrate preference and localisation. Alkbh5 is localised in the nucleus and demethylates m^6^A on DRACH motif. Whereas, FTO is localised both in the cytosol and the nucleus and acts on broad spectrum substrates^8^. Some studies have defined the role of FTO in neuron present in cytosol and dendrites, where it regulates local translation of mRNA ^9,10^. Thus, these studies provide basis for the FTO to study the role of activity mediated translation response. The m^6^A mark is recognised by a set of the proteins known as readers, which preferentially binds to the methylated RNA and mediate the downstream effector functions of m^6^A modification ^11^. The YTH domain-containing proteins YTHDF1-3, YTHDC1-2 are the most common m^6^A readers in mammals, and are proposed to play a role in determining the stability and translational efficiency of the bound transcripts ^12–14^.

Several recent studies have uncovered the role of m^6^A modification in the nervous system. The m^6^A modified transcripts are highly enriched in the brain ^5,15^, and their levels are developmentally regulated^16,17^. Studies involving knockdown and overexpression of m^6^A writers and erasers have demonstrated the role of m^6^A in the process of neurogenesis, axon morphogenesis, synapse morphogenesis and synaptic plasticity ^18,19^. Furthermore, studies have also shown the preferential synaptic localization of m^6^A marked transcripts, and experience dependent modulation of m^6^A levels^3^. The dynamic and reversible nature of m^6^A modifications makes it an excellent candidate for regulation protein synthesis upon synaptic activity. Surprisingly, despite the recent surge in studies focusing on synaptic m^6^A enrichment and experience dependent m^6^A modulation, the dynamic regulation of m^6^A in response to synaptic activity, and its relation to translation remains unexplored^20^. In our previous studies, we have shown the importance of translation regulation^21,22^ and its kinetics^23^ on synaptic stimulation. In this study, we probed the temporal dynamics of m^6^A in response to NMDAR stimulation. Briefly, we observe a robust increase in the total m^6^A levels, temporally coinciding with the global translation inhibition phase of NMDAR stimulation in cultured cortical neurons. This is facilitated by the reduction in the somato-dendritic expression of the m^6^A demethylase FTO along with the redistribution of m^6^A marked transcripts in the non-translating fractions of the polysome profile.

## Results and discussion

### NMDAR stimulation leads to increase in m^6^A levels on RNA which correlates with the global translation inhibition

In this study, we have used a well-established NMDAR stimulation paradigm to understand the change in m^6^A levels on RNA in response to synaptic activity in neurons ^24^. NMDAR is known to elicit a dynamic translational response in neurons, involving rapid and robust inhibition of global translation for a short term period (1-5 minutes), followed by the activation of global translation at delayed period (20 minutes) ^21,23,24^. At first, we validated our NMDAR stimulation paradigm by measuring the translation response in cultured rat cortical neurons upon NMDA treatment for 1, 5, and 20 minutes. We used the phosphorylation status of eEF2 to measure global translation response downstream of NMDAR stimulation. We measured the levels of p-eEF2 in cortical neurons using immunostaining analysis after 1, 5, and 20 minutes of treatment with 20 µM NMDA. We observed a robust, 3-fold increase in p-eEF2 levels upon 1-minute NMDAR stimulation (**Figure 1A and 1B**). By 5 minutes, the p-eEF2 levels had reduced compared to 1 minute time point; however, it remained significantly higher compared to unstimulated condition (**Figure 1A and 1B**). By 20 minutes, there was a maximum reduction in p-eEF2 levels, bringing it lower than the untreated condition (**Figure 1A and 1B)**. In addition, we have quantified the phosphorylation of eEF2 through western blotting and we observed a significant increase in p-eEF2 levels at 1 minute NMDAR stimulation **(Supplementary Figure 1A and 1B)**. These results are in accordance with the previous reports ^23,24^ which show a rapid translation inhibition followed by a delayed translation activation upon NMDAR stimulation.

**Figure 1.**
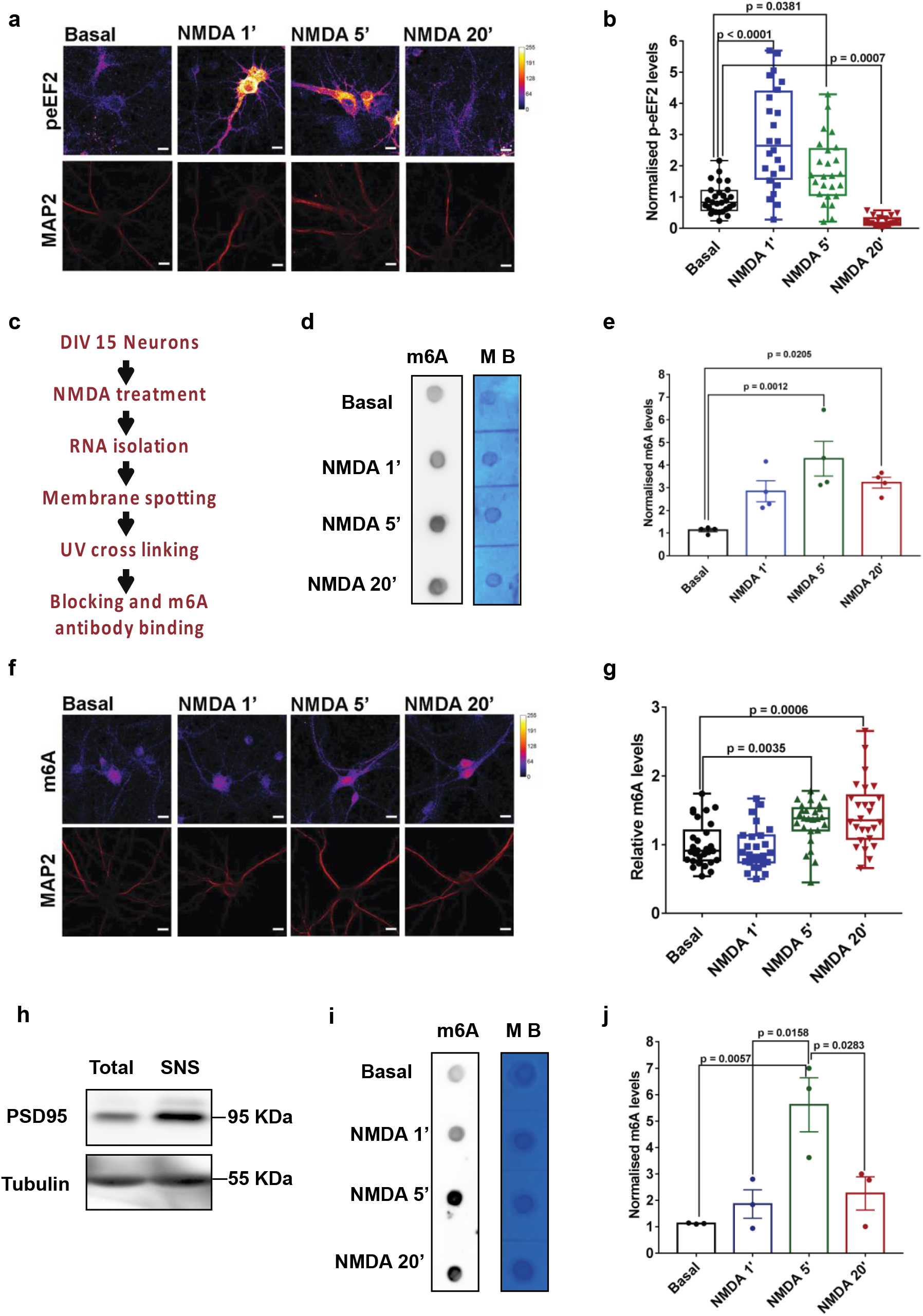
NMDAR stimulation leads to increase in m^6^A levels on RNA which correlates with the global translation inhibition response. **a-** Representative image showing p-eEF2 and MAP2 staining in DIV15 cultured cortical neurons treated with 20 µm NMDA for 1, 5 and 20 minutes. **b-** Quantification of p-eEF2 levels normalized to MAP2 in DIV15 cultured cortical neurons treated with 20 µm NMDA for 1, 5 and 20 minutes. Data represents mean +/-SEM, n > 22 neurons for all groups, from at least 3 independent neuronal cultures, Kruskal-Wallis test followed by Dunn’s multiple comparison test. **c-** Schematic depicting m^6^A dot blot procedure after 20 µm NMDA treatment in cultured cortical neurons. **d-** m^6^A immunoblot and methylene blue (MB) staining blot processed parallelly for the DIV15 cortical neurons treated with 20 µm NMDA for 1, 5 and 20 minutes. **e-** Quantification of the m^6^A immunoblot normalized to MB signal in 20 µm NMDA stimulated neurons. Data represents mean +/-SEM, N=4 independent neuronal cultures, one way ANOVA with Dunnett’s multiple comparison test **f-** Representative images showing m^6^A and MAP2 staining in DIV15 cultured cortical neurons treated with 20 µm NMDA for 1, 5 and 20 minutes. **g-** Quantification of m^6^A levels (normalized to MAP2) in DIV15 cultured cortical neurons treated with 20 µm NMDA for 1, 5 and 20 minutes. Data represents mean +/-SEM, n > 24 neurons for all groups, from at least 3 independent neuronal cultures, Kruskal-Wallis test followed by Dunn’s multiple comparison test. **h-** Representative immunoblots showing enrichment of PSD95 in synaptoneurosomes (SNS) samples. Sample indicating total lysate and SNS are stained against PSD95 and Tubulin as loading control. **i-** m^6^A immunoblot and control methylene blue stained blots processed parallelly for the synaptoneurosomes treated with NMDA (40 µM) for 1, 5 and 20 minutes. **j-** Quantification of m^6^A immunoblot normalized to MB signal for synaptoneurosome samples treated with 40 µm NMDA for 1, 5 and 20 minutes. Data represents mean +/-SEM, N=3 animals, One way ANOVA with Tukey’s multiple comparison test.

Next, to understand the NMDAR mediated changes in m^6^A kinetics, we used the RNA dot blot method to measure the total m^6^A levels upon 1, 5 and 20 minutes of NMDA treatment of cultured cortical neurons. Brief methodology and steps involved in RNA dot blotting are indicated in **Figure 1C**. Total RNA samples extracted from the DIV15 cultured neurons treated with NMDA (1,5 and 20 minutes) are indicated in the dot blot (**Figure 1D**). We have also shown that the observed m6A signal is majorly contributed by RNA and not by DNA **(Supplementary Figure 1C)**. Blots were probed for m^6^A and methylene blue (colorimetric read out) which was used as a loading control (**Figure 1D**). We observed a significant increase in total m^6^A levels on 5-minute and 20-minutes of NMDAR stimulation, while the 1-minute time point did not show any significant change as compared to the untreated condition **(Figure 1D and 1E)**. These results suggest that, at 5-minute NMDAR stimulation, the increased m^6^A levels correlate with the translation inhibition (increased phosphorylation of eEF2). Hence, we propose that measured m^6^A levels are inversely correlative to NMDAR translation response at the 5-minute time point. Interestingly, there is a time delay in the peaking of m^6^A levels (peaks at 5 minutes) as compared to eEF2 phosphorylation (peaks at 1 minute). This indicates that the change in m^6^A levels is likely to be downstream of the kinase activation upon NMDAR stimulation. Further, measured m^6^A levels at 20 minutes shows a decreasing trend compared to 5 minutes NMDAR stimulation, but remains significantly high in comparison to basal condition. These results suggest that the m^6^A mark on RNA can be used as a potential marker to understand the temporal profile of NMDAR-dependent translation response.

We further validated our dot-blot results by investigating the NMDAR mediated changes in neuronal m^6^A levels using immunostaining analysis. Similar to our previous experiment, we stimulated DIV15 cultured cortical neurons with 20 µM NMDA for 1, 5 and 20 minute time points followed by immunostaining with m^6^A and MAP2 antibodies. In accordance with our previous results from dot-blot analysis, our image quantification revealed that m^6^A levels were not altered at 1-minute, but significantly increased at 5-minute and 20-minute time points of NMDA treatment (**Figure 1F and 1G**).To validate the total neuronal m^6^A measurements in synaptic compartments, we used synaptoneurosomal preparations. Cortical synaptoneurosomes were prepared from P30 rats and stimulated them with 40 µM NMDA for 1, 5, and 20 minutes and investigated the changes in m^6^A levels using dot-blot analysis. The synaptoneurosomal preparation was validated by the enrichment of post-synaptic protein PSD95 using immunoblotting analysis (**Figure 1H**). We used dot blot to understand the m^6^A methylation pattern in synaptoneurosomes. Quantification of m^6^A dot blots from synaptoneurosome indicated that m^6^A levels increased upon NMDA treatment, peaking at 5-minute time point and showed a significant reduction at 20 minutes compared to 5 minutes (**Figure 1I and 1J**). This is consistent with our previous observations from cultured cortical neurons (**Figure 1B and 1E)**. Overall, it suggests that NMDAR mediated changes in the m^6^A signal is dynamic and inversely-correlated to translation response (initial increase and later decrease of m^6^A coincides with the initial translation inhibition followed by translation activation), both in the whole neuron and in the synaptic compartments.

### NMDAR stimulation induces changes in nuclear and cytosolic levels of m^6^A demethylase FTO

In the previous section, we showed the distinct temporal profile of m^6^A upon NMDAR stimulation. Next, we wanted to explore the possible mechanism behind NMDAR mediated changes in m^6^A levels. The dynamic changes in m^6^A levels is primarily determined by the action of designated methyltransferases and demethylases ^6^. The m^6^A methyltransferases primarily act in the nucleus in a co-transcriptional manner ^25^, while the m^6^A demethylases are known to function in both nuclear and cytoplasmic compartments ^9,26,27^. Since we observed the NMDAR induced changes in the m^6^A levels in the cell body as well as the synapto-dendritic compartments, we hypothesized that m^6^A demethylases are the primary determinant of NMDA-induced changes in m^6^A levels. Among the two widely known m^6^A demethylases, ALKBH5 and FTO, ALKBH5 is known to primarily localize in the neuronal nucleus and its levels are low in the adult brain ^26^. On the other hand, FTO is widely studied in neurons and is shown to be expressed in the nucleus, dendrites and dendritic spines of CA1 pyramidal neurons ^28^. Hence, we speculated that FTO is the primary driver of mediating NMDAR induced changes in m^6^A levels. To test this, we performed immunostaining to determine the nuclear and cyto-dendritic changes in the FTO levels on 1, 5, and 20 minutes treatment with 20 µM NMDA. In accordance to the previous studies ^28,29^, we observed that FTO was primarily localized in neuronal nucleus (**Figure 2A**). Notably, we were also able to detect the FTO staining in the cytosolic and dendritic compartments (**Figure 2A**). As we find high levels of FTO in the nucleus in comparison to the cyto-dendritic compartment, we imaged nuclear and cyto-dendritic FTO under different imaging parameters (**Supplementary Figure 2A and 2B)**. The cyto-dendritic quantification revealed that the levels of FTO significantly decreased upon 1-minute NMDAR stimulation and recovered to the basal levels by 5-minute time point (**Figure 2B**). We did not observe any significant difference in the FTO levels between untreated and 20-minute NMDA treated neurons (**Figure 2B)**. When a similar analysis was done for the nuclear FTO levels, we again found a significant reduction in the FTO levels in 1-minute NMDA treated neurons in comparison to the untreated neurons (**Figure 2C and 2D**). However, in contrast to the cyto-dendritic FTO levels, the nuclear FTO levels remained significantly low even after 5 minutes of NMDA treatment. On 20 minutes of NMDA treatment, the FTO levels had recovered and was significantly higher compared to the untreated condition (**Figure 2D**). To determine the total FTO levels we performed immunoblot on total protein lysate and we observed that significantly low amount of FTO at the 5 minutes **(Figure 2E and 2F)**. Thus, we observed a temporal delay between NMDAR induced reduction in FTO levels versus NMDAR mediated increase in m^6^A levels. A significant and consistent increase in m^6^A levels was observed by 5 minutes of NMDAR stimulation, whereas the reduction in the total FTO protein levels were consistently low at NMDA 5-minute time point.

**Figure 2.**
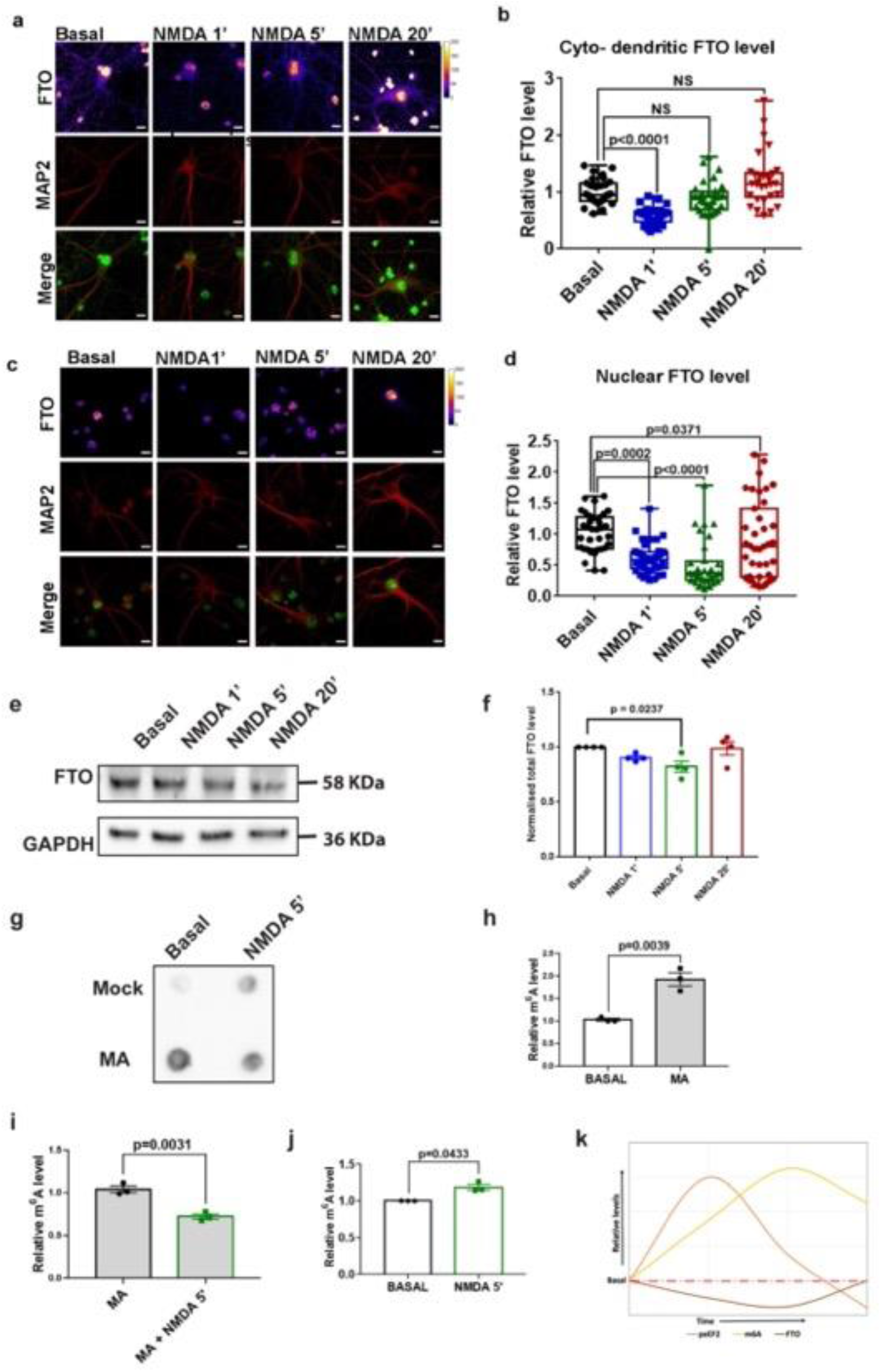
NMDA induces changes in nuclear and cytosolic levels of m^6^A demethylase FTO. **a-** Representative images showing FTO and MAP2 staining in DIV15 cultured cortical neurons treated with 20 µm NMDA for 1, 5 and 20 minutes. **b-** Quantification of cyto-dendritic FTO levels (normalized to MAP2) in DIV15 cultured cortical neurons treated with 20 µm NMDA for 1, 5 and 20 minutes. Data represents mean +/-SEM, n > 24 neurons for all groups, from at least 3 independent neuronal cultures, Kruskal-Wallis test followed by Dunn’s multiple comparison test. **c-** Representative images showing FTO and MAP2 staining in DIV15 cultured cortical neurons treated with 20 µm NMDA for 1, 5 and 20 minutes. **d-** Quantification of mean intensity of nuclear FTO levels in DIV15 cultured cortical neurons treated with 20 µm NMDA for 1, 5 and 20 minutes. Data represents mean +/-SEM, n > 24 neurons for all groups, from at least 3 independent neuronal cultures, Kruskal-Wallis test followed by Dunn’s multiple comparison test. **e-** FTO immunoblot and control GAPDH blots for the total FTO protein levels treated with NMDA (40 µM) for 1, 5 and 20 minutes. **f-** Quantification of FTO levels and plotted values are normalized to GAPDH. Data represent mean +/-SEM, N=4 independent experiments, One way ANOVA with Tukey’s multiple comparison test. **g-** m^6^A immunoblot for basal and NMDA 5 minute condition in presence of FTO inhibitor Meclofenamic acid (MA). **h-** Quantification of m^6^A signal comparing Basal and MA treatment for 2 hrs, N=3, paired T-test. **i-** Quantification of m^6^A signal in DIV15 cultured cortical neurons treated with 20 µm NMDA for 5 minutes with and without NMDA, N=3, paired T-test. **j-** Quantification of m^6^A signal in DIV15 cultured cortical neurons treated with 20 µm NMDA for 5 minutes and comparing Basal, N=3, paired T-test. **k-** Representation of temporal profiles of eEF2, total m^6^A, and total FTO levels.

To understand the importance of FTO in NMDA mediated changes of m^6^A levels, we used an FTO specific inhibitor Meclofenamic acid ^30^(MA) and compared the m^6^A levels at basal and NMDA stimulation conditions (**Figure 2G**). Treatment with MA (120 μmolar) for 2 hours caused a significant increase in m^6^A levels compared to basal condition validating the inhibition of FTO (**Figure 2H**). Further, 5-minute NMDAR stimulation in the presence of FTO inhibitor MA did cause significant changes in the m^6^A levels as compared to the mock (MA) treatment **(Figure 2I)**. Finally, as a control, we recaptured the increase in m6A signal upon 5-minute NMDAR treatment in the absence of MA **(Figure 2J)**. Thus, we show that FTO is a critical player mediating the dynamic changes of m^6^A levels upon NMDAR stimulation.

In the FTO imaging analysis, we observed differential dynamics of nuclear versus cyto-dendritic changes in FTO levels upon NMDAR stimulation. We speculate that this is primarily caused by redistribution of FTO between these compartments, along with the differential decay kinetics of nuclear and cyto-dendritic FTO. NMDAR induced changes in FTO levels could be attributed to the transcription, translation as well as degradation pathways. We summarise our findings in the representative graph shown in **Figure 2K**. The m^6^A readout follows a similar trend as eEF2 phosphorylation, but with delay in reaching the peak (**Figure 2K**). The reduction of the total FTO levels show a good correlation with increase in total m^6^A levels (**Figure 2K**). It is likely that the reduction in the FTO levels on 1 minute and 5 minutes of NMDAR stimulation is caused by ubiquitin-mediated degradation, as NMDAR activation is reported to cause widespread degradation at acute time points ^23^. Further, NMDA treatment is also known to cause a delayed translation activation response, providing a possible explanation for the increase in FTO levels on 20 minutes treatment ^23,24^.

### NMDAR mediated increase in m^6^A levels is accompanied with the shift of m^6^A marked RNA from polysome to non-polysome fractions

Our previous experiments clearly demonstrated the increase in m^6^A levels upon 5 minutes of NMDA treatment. From our previous observation, it is known that NMDAR stimulation elicits an overall translation inhibition response at 5-minute time point ^24^. We wanted to understand if the NMDAR mediated increase in m^6^A levels drives the translation repression of m^6^A marked RNAs. To investigate this, we used polysome profiling technique to monitor the distribution of m^6^A marked RNAs in ribosomal/polysomal pool versus non-ribosomal pool on NMDA treatment. Briefly, the DIV15 cultured cortical neurons were treated with 20 µM NMDA for 1, 5 and 20 minutes and the lysates were separated on 15-45% linear sucrose gradient. The steps involved in the sample treatment, polysome profiling and pooling strategy are depicted in **Figure 3A**. A representative profile (A254) is shown in **Figure 3B**. Additionally, we show the 18S rRNA distribution in the translating and non-translating fractions in the basal condition (Supplementary Figure 3A).

**Figure 3.**
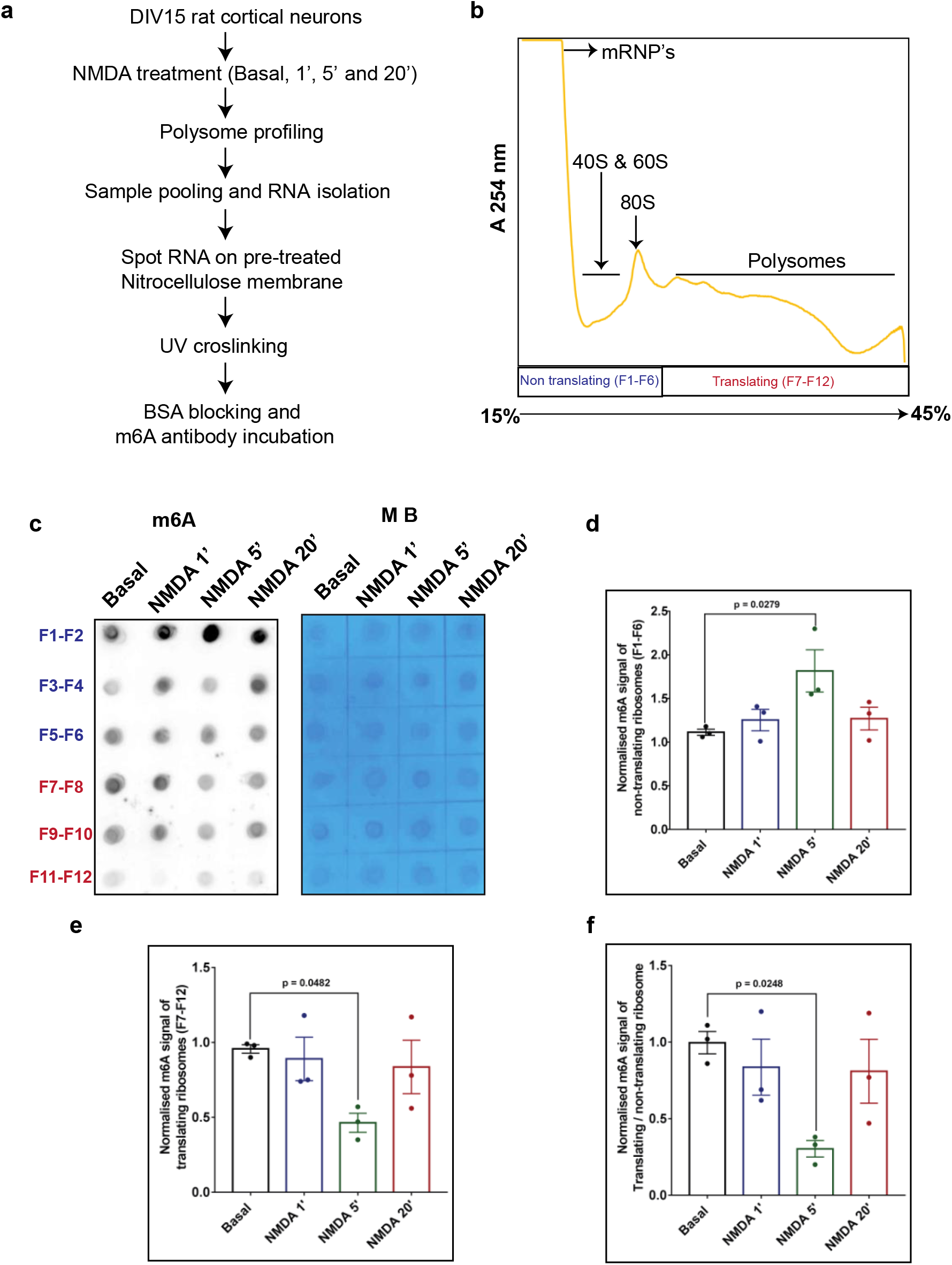
NMDAR mediated increase in m^6^A levels is accompanied with shift of m^6^A marked RNA from polysome to non-polysome fractions. **a-** Schematic representing detailed methodologies for sample preparation, polysome profiling and dot blot to analyze polysome distribution of m^6^A on 20 µm NMDA treatment. **b-** Representative polysome profile or absorbance profile (254 nm) run on 15 to 45 % sucrose gradient for the DIV15 rat cortical neurons and sample pooling strategy indicating non-translating pool(in purple) and translating pool (in red). **c-** m^6^A dot blot showing pooled polysome fractions on vertical lane and NMDA treatment time points (0, 1, 5 and 20 minutes) on horizontal lane. Control methylene blue (MB) blots showing the pooled polysome fractions on vertical lane (processed parallelly) and NMDA stimulation time points on the horizontal lane. **d-** Quantification of m^6^A signal in non-translating pool of polysome samples obtained from NMDA treated cortical neurons. Data represents mean +/-SEM, N=3 independent cultures, One way ANOVA with Dunnett’s multiple comparison test. **e-** Quantification of m^6^A signal in translating pool of polysome samples obtained from NMDA treated cortical neurons. Data represents mean +/-SEM, N=3 independent cultures, One way ANOVA with Dunnett’s multiple comparison test. Quantification of the ratio of m^6^A signal from translating pool to non-translating pool of polysome samples obtained from NMDA treated cortical neurons. Data represents mean +/-SEM, N=3 independent cultures, One way ANOVA with Dunnett’s multiple comparison test.

As shown in schematic **Figure 3B**, we have labelled the pooled samples from F1-6 as non-translating and F7-12 as translating pool, indicated in purple and red colour respectively. An equal volume of each fraction was taken for RNA isolation and subjected to dot blot with m^6^A antibody and methylene blue staining to generate the calorimetric signal (loading control) (**Figure 3C**). We observed a uniform staining in methylene blue treated samples, indicating approximately equal loading of RNA on the membrane. We observed a significant increase in m^6^A levels in the non-translating pool upon 5-minutes NMDA treatment (**Figure 3D**). In contrast, quantification of the translating pool of ribosomes showed a decrease in m^6^A signal at 5 minutes of NMDA treatment (**Figure 3E**). Further, quantification of the ratio of m^6^A levels in the translating to non-translating pool showed a significant decrease of the m^6^A levels at 5-minute NMDAR stimulation compared to basal condition (**Figure 3F**). This decrease in m^6^A levels in the translating fractions suggests that the m^6^A marked pool of ribosomes and mRNAs associated with them have shifted towards the non-translating pool, indicative of translation inhibition. Altogether this data suggests that the NMDAR induced increase in overall m^6^A signal is accompanied by the redistribution of m^6^A marked RNA from translating to non–translating fractions. This shift in signal from translating to non-translating pool could potentially be contributed by both rRNA and mRNA and we are yet to identify the factors driving this shift. In contrast, we observe that the m^6^A levels does not alter significantly at 1 and 20 minutes of NMDA treatment. Further, since the m^6^A mark on these RNAs could be potentially removed by demethylases, these RNAs could shift back to the translating pool. Another interesting possibility is that the pool of RNAs present in m^6^A marked inhibitory complex could be targeted for enzymatic degradation ^31,32^. Thus, from our observations, we conclude that m^6^A signal dynamically distributed across the polysome fractions upon NMDAR stimulation.

Apart from mRNA m^6^A modification, there are reports suggesting m^6^A role in non-coding RNA modification and known to affect gene expression ^33,34^. Non-coding RNA like microRNA, tRNA, rRNA and lncRNA are m^6^A methylated and their changes are implied in diseases such as cancer ^34,35^. In case of microRNA, presence of m^6^A is known to reduce the duplex stability between the 3’UTR and miRNA seed region ^36,34^. Other prime example is from the rRNA, where 18S and 28S rRNA carry one m^6^A mark each which is shown to regulate protein synthesis ^37^. While our interpretation is mainly focused on m^6^A modifications on mRNA, we cannot rule out the changes in m^6^A mark on other RNAs contributing to our results.

Overall, we have shown that m^6^A levels change dynamically upon NMDAR stimulation. At 5-minute NMDAR stimulation, we observe an increase in m^6^A levels which is correlated with a phase of translation inhibition. Correspondingly, m^6^A signal is also shifted from the translating fractions towards the non-translating pool at 5-minute NMDAR stimulation; further supporting the correlation with translation inhibition. Interestingly, the levels of m^6^A demethylase FTO is decreased at 5-minute NMDAR stimulation which is responsible for the increase in m^6^A levels. Inhibition of FTO prevents the dynamic changes of m^6^A levels upon NMDAR stimulation indicating that FTO is a key player in this regulation.

## Methods

### Ethics Statement

The study was carried out in compliance with the ARRIVE guidelines. We performed all the animal work in accordance to the guidelines approved by the Institutional Animal Ethics Committee (IAEC) and the Institutional Biosafety Committee (IBSC), InStem, Bangalore, India. All experiments were performed with a minimum of three independent neuronal cultures. All our experiments were performed with cultured neurons and synaptoneurosomes preparation derived from Sprague Dawley (SD) rats. Rat colonies were maintained at 14 hour/10 hour light/dark cycle, 20 - 22 °C temperature, 50 – 60% relative humidity. The rooms harbouring the colonies were supplied with 0.3 µm HEPA-filtered air. Rats were freely fed with food and water.

### Primary neuronal culture and Inhibitor treatment

Primary neuronal cultures were prepared cortices of embryonic day 18 (E18) Spargue-Dawley rats following previously established protocols ^38,39^. Briefly, the dissociated cortices were trypsinised with 0.25 % trypsin, followed by washes with sterile HBSS and resuspension in Minimum Essential Media (MEM, 10095080, Thermo Fisher Scientific) containing 10 % FBS (F2442, Sigma-Aldrich). The cells were then counted and plated at a density of 40000 cells/cm^2^ on poly-L-lysine (P2636, Sigma-Aldrich) (0.2 mg/ml in borate buffer, pH 8.5) coated dishes. After 3 hours of plating, the media was changed to neurobasal (21103049, Thermo Fisher Scientific) supplemented with B27 (17504044, Thermo Fisher Scientific) and Glutamax (35050-061, Life Technologies). Neurons were cultured for two weeks at 37 °C in a 5% CO_2_ incubator, and supplemented with neurobasal after every five days. On DIV15 the neurons were treated with 20 µM NMDA for 1, 5 and 20 minutes time points and were processed for downstream experiments as per requirement. FTO Inhibitor (Maclofenamic acid) treatment was done at DIV15 stage at 120 μmolar MA for 24 hrs and followed by NMDA treatment for 5 minutes. After treatment with NMDA samples were separated for protein and RNA work.

### Immunostaining

NMDA treated DIV15 cortical neurons were fixed with 4 % Paraformaldehyde (PFA) for 20 minutes at room temperature and processed for subsequent immunostaining analysis. Briefly, the fixed neurons were washed with PBS to remove the traces of PFA, followed by 10 minutes permeabilization with TBS_50_T (0.3 %) (50 mM Tris-HCl (pH 7.4), 150 mM NaCl, 0.3% Triton X-100) and 1 hour blocking at room temperature with the blocking buffer (TBS_50_T (0.1 %), 2 % BSA, 2 % FBS). Subsequently, the neurons were incubated with required primary antibodies for two hours at room temperature. Afterwards, the neurons were washed with washes with TBS_50_ T (0.1 %), and incubated with the required secondary antibodies for one hour at room temperature. After final washes, the coverslip with neurons were mounted on slide using Mowiol 4-88 mounting media. All the Images were captured on FV3000 confocal microscope (Olympus) at 60X, NA 1.4, oil immersion objective, pinhole set at one airy unit. The imaging parameters were kept constant across different time points in an experiment.

### Western blot

DIV15 cortical neurons treated with 20 μM NMDA for different time periods were lysed in lysis buffer containing 50 mM Tris (pH 7.4), 150 mM NaCl, 5 mM MgCl_2_, 1 % Triton X-100, supplemented with EDTA-free protease inhibitor complex (Cat.no-S8830, Sigma) and phosphatase inhibitor cocktail (Cat.no-04906837001, Roche). The cells were subsequently centrifuged at 16000 RCF for 30 min at 4 ^°^C and the obtained lysates were resuspended in laemmli buffer and were heat denatured at 95 ^°^C for 5 min. The samples were stored at -20 ^°^C until further use. 10 % PAGE gel was prepared and 15 μL of sample was loaded onto each well and 1.5 hr transfer was done at 4 ^°^C. Blots were stained for the control Ponceaus staining to verify the transfer and after washing blot was blocked for 1 hr at room temperature in TBST with 5 % BSA. For primary staining we have used total eEF2 (Cat.no-2331S, CST), peEF2 (Cat.no-2332S, CST), FTO (Cat.no – 45980, CST) and GAPDH (Cat.no-2118S, CST) as a loading control, secondary antibody (Cat.no-A0545, Sigma-Aldrich) with HRP conjugation was used and Clarity western ECL (Bio-Rad) was used to develop and imaged in the GE Amersham imager 600. For the all the replicative immunoblots of FTO and eEF2 blots were cut at their corresponding molecular weight makers and probed later with respective antibodies. This method of blot imaging was done for simultaneous imaging of blots and to reduce the variance in the assay.

### Polysome profiling

The DIV15 rat cortical neurons were stimulated with NMDA (20 μM) for 1, 5 and 20 minutes. After treatment cells were lysed in lysis buffer (20 mM Tris-HCl, 100 mM KCl, 5 mM MgCl_2_, 1% NP40, 1mM dTT, 1X Protease inhibitor cocktail, RNAse inhibitor, 0.1 mg/mL cylohexamide and 1X Phosphotase inhibitor) and centrifuged at 20K X g at 4 ^°^C for 20 min. The supernatant was loaded onto 15-45% linear sucrose gradient prepared in buffer (20 mM Tris-HCl, 100 mM KCl, 5 mM MgCl_2_, 0.1 mg/mL Cycloheximide). The gradient was loaded with cell lysate and centrifuged at 39K RPM at 4°C for 90 min. Gradient fractions were collect using Brandel fractionation collector instrument and equal volume of each pooled fractions were processed for RNA isolation and dot blot.

### RNA isolation and Dot blot

The DIV 15 rat cortical neurons and synaptoneurosomal, RNA was isolated using standard TRIzol RNA extraction method (Thermo Fisher Scientific cat.no – 15596018). Isolated RNA was finally resuspended in milliQ water and its concentration was measured using nanodrop and equal an concentration of RNA was used for dot blot analysis. For the dot blot, the Nitrocellulose membrane (Cat.no-10600002, Sigma-Aldrich) was cut according to the requirement and rinsed first with milliQ water followed with 20X SSC buffer (Cat.no-AM9763, Thermo Fisher Scientific) and air dried. Extracted RNA was diluted to 250 ng in the RNA dilution buffer (6X SSC buffer and 7.5% para-formaldehyde) and heated to 65 ^°^C for 5 minutes and kept on ice for 5 min. RNA was spotted on to nitrocellulose membrane and UV crosslinked. The membrane was blocked in TBST with 5% BSA for 1 hour at room temperature. Further, membrane was incubated overnight at 4 ^°^C with 1:1000 dilution of m^6^A antibody (Cat.no-202 111, Synaptic systems) Subsequently, the membrane was washed three times in TBST for 10 min of interval. Anti-rabbit-HRP secondary antibody was incubated at 1:10,000 dilution for 1 hour at room temperature.

### Methylene-blue staining

RNA was extracted from the respective samples using the standard Trizol-LS protocol (Cat.no-10296010, Thermo Fisher Scientific). The Nitrocellulose membrane (Cat.no-10600002, Thermo Fisher Scientific) was cut according to the requirement and rinsed with milli Q water and 10X SSC. Afterwards the membrane was airdried until the loading of samples. The samples were prepared by diluting the extracted RNA to a final concentration of 250 ng in the RNA dilution buffer followed by heating of diluted samples at 65 ^°^C for 5 minutes. Afterwards the samples were incubated on ice for 5 minutes. The diluted RNA sample were spotted on the activated membrane and crosslinked using UV cross linker. After crosslinking, the membrane was incubated with the methylene blue staining solution (0.3 M sodium acetate and 0.03 % methylene blue) for 5 minutes, followed by washes with distilled water to remove the background signal. The processed membrane was then imaged using Image Quant (LAS 4000 / Amersham imager 600).

### Quantitative PCR

Isolated RNA from the NMDA treated sample was processed for cDNA synthesis with reverse transcriptase and without reverse transcriptase. 200 ng of RNA was taken and cDNA synthesis was done with random hexamer and while doing the reverse transcription enzyme (M-MLV cat.no 28025013, Invitrogen) was excluded (minus reverse transcription) and included (plus reverse transcription) to the master mix and cDNA synthesis was done according to manufacturer protocol.Quantification of PSD95, PTEN and actin was measured by qPCR using TAKARA SYBR green (Cat.no – RR82WR).

### Synapto-neurosome preparation

Rat cortical synaptoneurosomes were prepared using the filtration method from Sprague Dawley (SD) rat^21,22^. Briefly, the dissected cortices were resuspended in the synaptoneurosome buffer (118 mM NaCl, 5 mM KCl, 1.2 mM MgSO_4_, 2.5 mM CaCl_2_, 1.53 mM KH_2_PO_4_, 212.7 mM Glucose, 1X Protease Inhibitor Cocktail, pH 7.5) followed by homogenization on ice. The obtained homogenate was filtered by passing through three 100 µM nylon filters (NY1H02500, Merck Millipore) and one 11 µM nylon filter (NY1102500, Merck Millipore). The filtrate was then centrifuged at 1500 RCF for 15 minutes at 4 ^°^C. The pellet obtained was resuspended in 2 mL synaptoneurosome buffer and used for NMDA treatment (40 µM) for different time points. After NMDA treatment, the resuspended synaptoneurosomes were subjected to a brief spin and the obtained pellets were resuspended in lysis buffer (50 mM Tris pH 7.4, 150 mM NaCl, 5 mM MgCl_2_, 1% Triton X-100, supplemented with EDTA-free protease inhibitor complex and phosphatase inhibitor cocktail) and centrifuged at 16000 RCF for 30 minutes at 4 ^°^C. The obtained lysates were used for western blotting and RNA isolation as per previously described protocol.

### Statistical analysis

Statistical analysis was performed using GraphPad prism software version 7.0.0. Prior to the calculation of differences between groups, the data distribution was tested for normality using Kolmogorov Shapiro Smirnov goodness-of-fit test and depending on the distribution, either parametric or non-parametric tests were used to calculate the statistical significance. For groups with less than 5 data points, data distribution was assumed to be normal. Multiple group comparisons were made using one-way ANOVA followed by Bonferroni’s multiple comparison test for normally distributed data and Kruskal-wallis followed by Dunn’s multiple comparison test. All the tests were doing by using GraphPad Prism version 7.0.0 for Windows, GraphPad Software, San Dieg, California USA, www.Graphpad.com.

## Acknowledgements

We thank all present and past Ravi Muddashetty lab members for the suggestion and support throughout this work.

## Authors contribution

NKCG and RM – Conceptualise the question, NKCG, BN and RM designed experiment; NKCG, BN, VJ – Performed experiments; SR-Provided the resources; NKCG, BN and RM Analysed data and wrote manuscript.

## Conflict of Interest

The authors declare that they have no conflict of interest.

## Notes

### Competing Interest Statement

The authors have declared no competing interest.

