## Supplementary Figures for "NMDAR mediated dynamic changes in m^6^A inversely correlates with neuronal translation"

### Supplementary Figure 1:

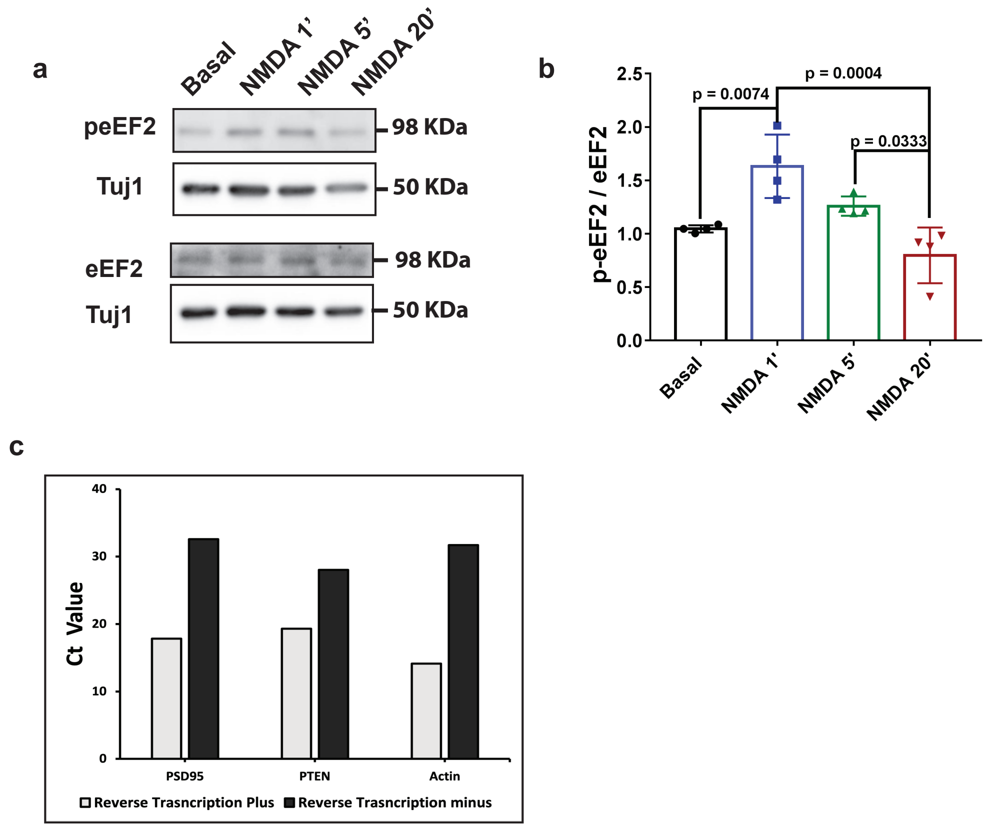


**Supplementary Figure 1: NMDAR stimulation leads to increase in m$^6$A levels on RNA which correlates with the global translation inhibition response**

**a-** Representative immunoblots showing p-eEF2 and eEF2 levels in DIV15 cultured cortical neurons treated with 20 µm NMDA for 1, 5 and 20 minutes and loading control indicated by Tuj1.

**b-** Quantification of p-eEF2 levels normalized to Tuj1 in DIV15 cultured cortical neurons treated with 20 µm NMDA for 1, 5 and 20 minutes. Data represents mean +/- SEM, n =4 from independent neuronal cultures, One-way ANOVA (p=0.0006) followed by Tukey's multiple comparison test.

**c-** Quantitative PCR to access the DNA contamination in RNA isolated from basal condition for mRNA candidates PSD95, PTEN and Actin plotted data points are the Ct values.

#### Supplementary Figure 2:

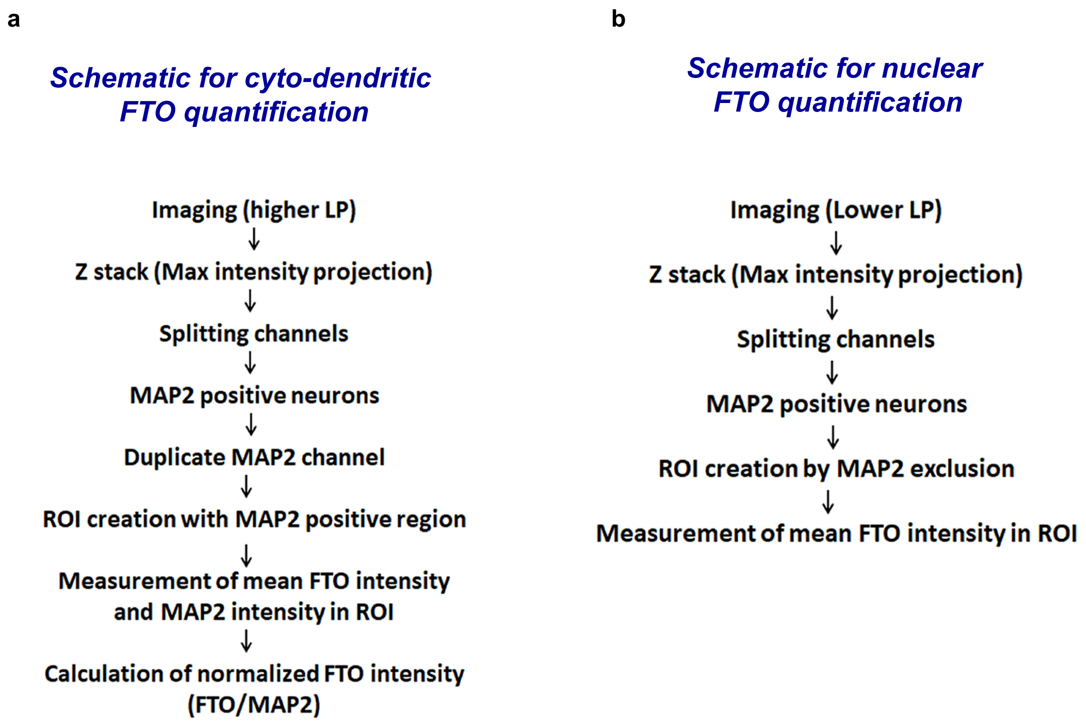


**Supplementary Figure 2: NMDA induces changes in nuclear and cytosolic levels of $m^6A$ demethylase FTO**

- **a-** Schematic depicting procedure used for quantification of cyto-dendritic FTO levels.

- **b-** Schematic depicting procedure used for quantification of nuclear FTO levels.

#### Supplementary Figure 3:

a

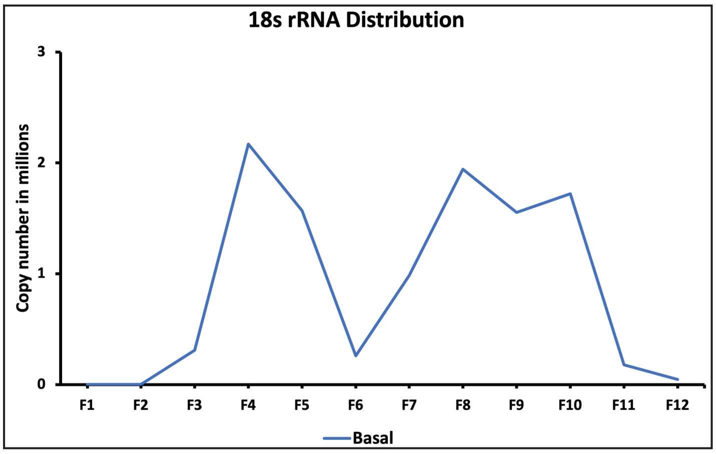


**Supplementary Figure 3: NMDAR mediated increase in m$^6$A levels is accompanied with shift of m$^6$A marked RNA from polysome to non-polysome fractions.**

**a-** Quantification of 18S rRNA by qPCR method and line graph indicate distribution across the polysome pools
